# An open dataset of cerebral tau deposition in young healthy adults based on [^18^F]MK6240 positron emission tomography

**DOI:** 10.1101/2025.06.13.655622

**Authors:** Jack Lam, Raúl Rodriguez-Cruces, Thaera Arafat, Jessica Royer, Judy Chen, Arielle Dascal, Ella Sahlas, Raluca Pana, Robert Hopewell, Chris Hung-Hsin Hsiao, Gassan Massarweh, Jean-Paul Soucy, Pedro Rosa-Neto, Sylvia Villeneuve, Lorenzo Caciagli, Matthias Koepp, Andrea Bernasconi, Neda Bernasconi, Boris Bernhardt

## Abstract

Tauopathies are pathologies wherein phosphorylated insoluble tau aggregates in neurons, leading to dysfunction and degeneration. Positron emission tomography (PET) enables measurement of *in vivo* tau, with second-generation radiotracers such as [^18^F]MK6240 showing high tau affinity with minimal off-target binding. While tauopathies are commonly linked to age-related neurodegenerative diseases, notably Alzheimer’s disease (AD), evidence suggests pathophysiological cascades may begin long before clinical onset. Increasingly, tau is recognized in pathologies affecting younger individuals, including autosomal dominant AD, Niemann-Pick disease type C, chronic traumatic encephalopathy, and epilepsy, thus highlighting the importance of normative data in non-geriatric populations. Here, we present a dataset of 33 young to middle-age healthy adults (mean age 34.0±10.4 years, 12 female) with [^18^F]MK6240 PET data and T1w magnetic resonance imaging. Longitudinal data are also available in a subset of 9 participants with a minimum follow-up time of 1 year. Our dataset aims to support imaging biomarker studies on younger individuals potentially at risk for AD and to advance work in tauopathies affecting non-geriatric populations generally excluded from neurodegeneration studies.

## Background

Tau is a protein encoded by the microtubule-associated protein tau (MAPT) gene located on chromosome 17q21 and is primarily involved in microtubule assembly and stabilization^1^ Hyperphosphorylation of tau reduces its binding affinity to microtubules and promotes aggregation into insoluble intracellular filaments, leading to neural toxicity and neurodegeneration.^2,3^ Tau aggregates in the central nervous system are the main characteristic of a group of diseases called tauopathies, which include pathologies such as Alzheimer’s disease (AD), Pick’s disease, frontotemporal dementias, progressive supranuclear palsy, and corticobasal degeneration.^4^ Tauopathies are not exclusive to diseases associated with aging, however. Autosomal dominant AD with underlying mutations in presenilin 1, presenilin 2, or amyloid precursor protein genes may present as early as the third decade of life.^5^ Niemann-Pick disease type C, which manifests abnormal tau protein similar to that in AD, may present at any age, however, prevalence peaks at age 19-30.^6,7^ Chronic traumatic encephalopathy (CTE) presents at a wide age range depending on the nature of trauma and shows a spectrum of hyperphosphorylated tau pathology.^8,9^ Additionally, increasing work looking into the bidirectional relationship between AD and seizures has sparked interest in the role of tau in epilepsy.^10^ Studies on surgically resected specimens and *post mortem* tissue show evidence of increased tau even in young to middle-aged adults with temporal lobe epilepsy as well as other developmental and acquired epilepsies.^11–14^ As our understanding of various tauopathies advances, there is an increasing need for non-invasive tools to study tau *in vivo*, which will provide important insights into disease progression, offer prognostic information, and support the development of therapeutic interventions targeting tau.

Over the past decade, numerous positron emission tomography (PET) radiopharmaceuticals have been developed to measure tau *in vivo*.^15^ The first-generation of tau PET radiotracers has been used extensively in research and includes [^18^F]THK5317, [^18^F]THK5351, [^11^C]PBB3, and [^18^F]AV1451 (also known as [^18^F]flortaucipir).^16^ Among these, [^18^F]flortaucipir was the first to be approved for clinical use by the Food and Drug Administration (FDA). [^18^F]flortaucipir showed high affinity and specificity for tau and correlated well with the spatial distribution of neurofibrillary tangles described by Braak staging.^17–19^ Despite its widespread adoption and validation for the diagnosis of AD, its utility in non-AD tauopathies is limited. Autopsy studies have shown negligible to poor flortaucipir binding in other tauopathies due to differences in tau isoforms.^20–22^ Off-target binding to melanin-containing structures, such as the substantia nigra, and calcified structures, such as the choroid plexus has also been noted in flortaucipir studies.^15,23^ Additionally, flortaucipir shows a relatively low effect size in non-AD tauopathies and has poor correlation with disease severity.^24^

Research is ongoing into second-generation radiotracers with the aim of developing tracers with broader applicability, improved specificity, and reduced off target binding.^15,25^ [^18^F]MK6240 is one promising second-generation tau PET tracer that shows high cell permeability, high affinity for neurofibrillary tangles, and poor binding to amyloid plaques.^26–28^ While there is noted meningeal and neuromelanin binding as well as weaker binding to intraparenchymal hemorrhage, [^18^F]MK6240 is not affected by the off-target binding in basal ganglia and choroid plexus that is seen with flortaucipir.^27,29^ Studies have demonstrated that [^18^F]MK6240 uptake in AD patients shows patterns consistent with the known distributions of tau deposition^30,31^ and may discriminate cognitively normal AD subjects from AD dementia patients,^30,32^ correlating with cognitive deficits.^33,34^ Moreover, similar to [^18^F]flortaucipir, increased [^18^F]MK6240 uptake in AD patients correlated with Braak staging, demonstrating its ability to detect early accumulation of neurofibrillary tangles and to follow its progression.^30,33,35^

In keeping with the open science mission of the Montreal Neurological Institute (MNI),^36^ we present a unique openly available dataset of healthy, young to middle-aged adult individuals with [^18^F]MK6240 PET and T1-weighted structural MRI. We include raw 3D PET and MRI volumes, MNI152 transformation matrices, cerebellar grey matter and composite^37^ (whole cerebellum, brainstem, and eroded subcortical white matter) reference regions, and partial volume corrected standard uptake maps based on these reference regions, as well as their respective surfaces in fsLR32k surface space. As brain changes may predate clinical AD symptoms by years,^38^ this provided data can help support studies on younger populations at risk for later development of AD in order to better understand pathophysiological pathways and identify potential interventions. Additionally, we present a subset of participants who have a repeat [^18^F]MK6240 PET scan with a follow-up time of at least one year, which will be useful to researchers working on presumably non-AD tauopathies affecting young to middle-aged adult individuals who have traditionally been underrepresented in research on tau.

## Methods

### Participants

Thirty-three healthy controls (mean age±standard deviation (SD) 34.0±10.4, 12 female) were recruited from the local Montreal area via advertisement. All participants signed a research ethics board (REB) approved informed consent form prior to participation (2018-4148, 2018-3469), including consent to share anonymized data in an openly shared repository. All participants denied a history of traumatic brain injury or neurological or psychiatric illness and had a Montreal Cognitive Assessment score of ≥26 (mean score±SD 28.1±1.7).

### MRI acquisition

An overall schematic of the imaging processing methods is presented in **Figure 1**. Protocolized structural MRI scans were carried out at the Brain Imaging Centre of the Montreal Neurological Institute using a 3T Siemens Magnetom Prisma-Fit equipped with a 64-channel head coil. The full scanning protocol can be found in Royer et al. 2022.^39^ T1-weighted (T1w) imaging was obtained using a 3D magnetization-prepared rapid gradient-echo sequence (MPRAGE; 0.8mm isotropic voxels, TR = 3200ms, TE = 3.14ms, TI = 900 ms, flip angle = 9 degrees).

**Figure 1.**
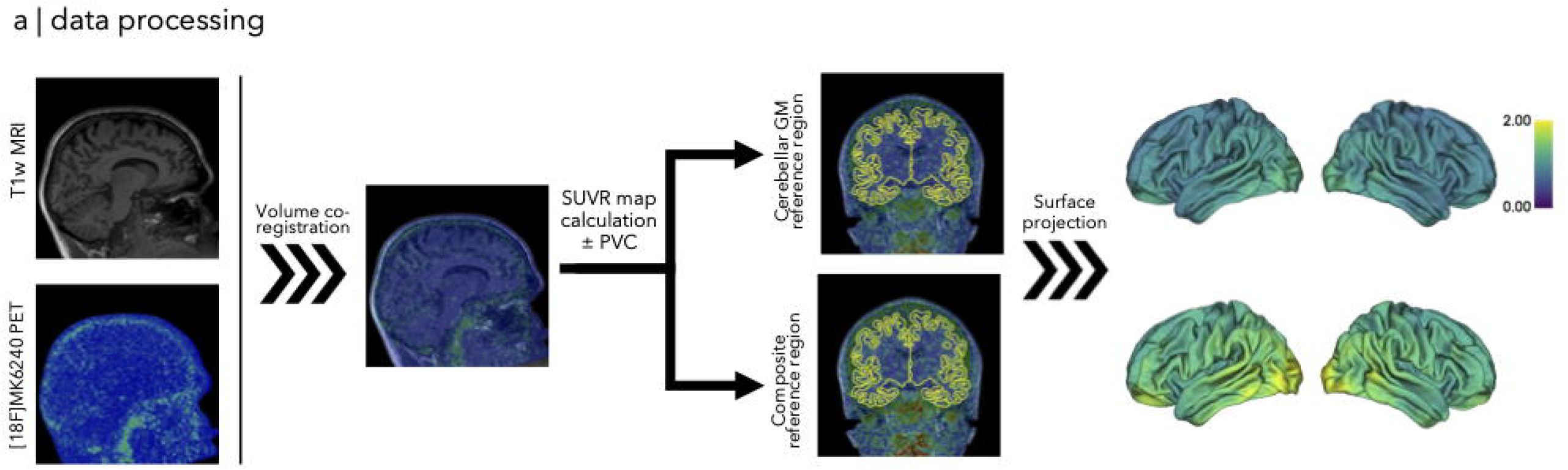
Overview of the data processing pipeline. The T1w MRI and 3D [^18^F]MK6240 PET were linearly coregistered and standard uptake value ratio (SUVR) maps were calculated using either the cerebellar grey matter or composite ROI reference region. Partial volume correction was done using PETPVC and both corrected and uncorrected SUVR maps were projected to fsLR32k surface space and smoothed to 10mm full-width at half-maximum. Abbreviations: SUVR, standard uptake value ratio; PVC, partial volume correction; GM, grey matter.

T1w MRI were processed using micapipe v0.2.3 (https://micapipe.readthedocs.io/), an openly available robust pipeline for multimodal MRI analysis,^40^ as well as FreeSurfer v6.0 (https://surfer.nmr.mgh.harvard.edu/) for cortical surface reconstruction and segmentation. All surfaces and segmentations were manually inspected and corrected. Each T1w MRI was deobliqued and reoriented to LPI orientation (left-right, posterior-anterior, inferior-superior).

### [^18^F]MK6240 PET acquisition and image processing

All participants underwent [^18^F]MK6240 during a separate session from the MRI. A subset of 9 participants underwent a second PET scan with the same acquisition protocol (mean interval 826±486.6 days, minimum 372 days, maximum 1797 days). [^18^F]MK6240 was synthesized in the PET unit of the McConnell Brain Imaging Centre according to previously published protocols.^41^ PET scans were acquired using a brain-dedicated Siemens high-resolution research tomograph (HRRT; Erlangen, Germany) scanner with a resolution of 2.4mm full-width at half-maximum (FWHM). A bolus of [^18^F]MK6240 (mean dose 240±18 MBq) was injected intravenously and images were acquired 90-110min post-injection (4 frames, 300s each). An ordered subset expectation maximization (OSEM) algorithm was used to reconstruct the images. A 6min transmission scan with a rotating Cesium-137 source was used for attenuation correction.

Images were corrected for dead time, decay, and random and scattered coincidences. Motion correction was performed using a coregistration-based frame realignment method for dynamic PET volumes that also compensates for emission-transmission mismatches.^42^ The dynamic [^18^F]MK6240 images were co-registered and averaged together to a single 3D volume.

The [^18^F]MK6240 PET images were linearly co-registered to the T1w MRI using Advanced Normalization Tools (ANTs).^43^ Cerebellar grey matter^30^ and composite (whole cerebellum, brainstem, and eroded subcortical white matter)^37^ reference regions were derived from the aparc atlas from FreeSurfer outputs. The average uptake in these reference regions was then used to calculate the respective standard uptake value ratio (SUVR) maps. SUVR is a widely used metric in [^18^F]MK6240 PET studies and previous studies have shown SUVR measured 90 minutes post radiotracer injection provides a robust estimate of neurofibrillary tangle load.^30^ Partial volume correction was undertaken using Müller-Gärtner correction with the PETPVC toolbox (https://github.com/UCL/PETPVC) using a point spread function (PSF) of 2.4mm based on the FWHM of the HRRT scanner.^44^ Corrected and uncorrected SUVR maps were sampled along each participants native cortical surface space and then resampled to fsLR32K standard space.^45^ PET data were spatially smoothed on the surface with a 10mm FWHM Gaussian kernel using Workbench Command.^46^

## Data Records

The provided raw data and derivatives are compliant with the Brain Imaging Directory Structure (BIDS) format.^47^ All data are available on the Open Science Framework (OSF; https://doi.org/10.17605/OSF.IO/ZNT9D).^48^

### Raw data

The raw T1w MRI (defaced for anonymity), [^18^F]MK6240 PET (4D image with 4 frames), and their respective .json files are provided in */sub-#/ses-01/anat* and */sub-#/ses-01/pet* (**Figure 2A**). Since only one MRI session was conducted, only the session-2 PET scan (and no MRI) is provided in */sub-#/ses-02*. Both session-1 and session-2 PET scans were registered to the session-1 MRI.

**Figure 2.**
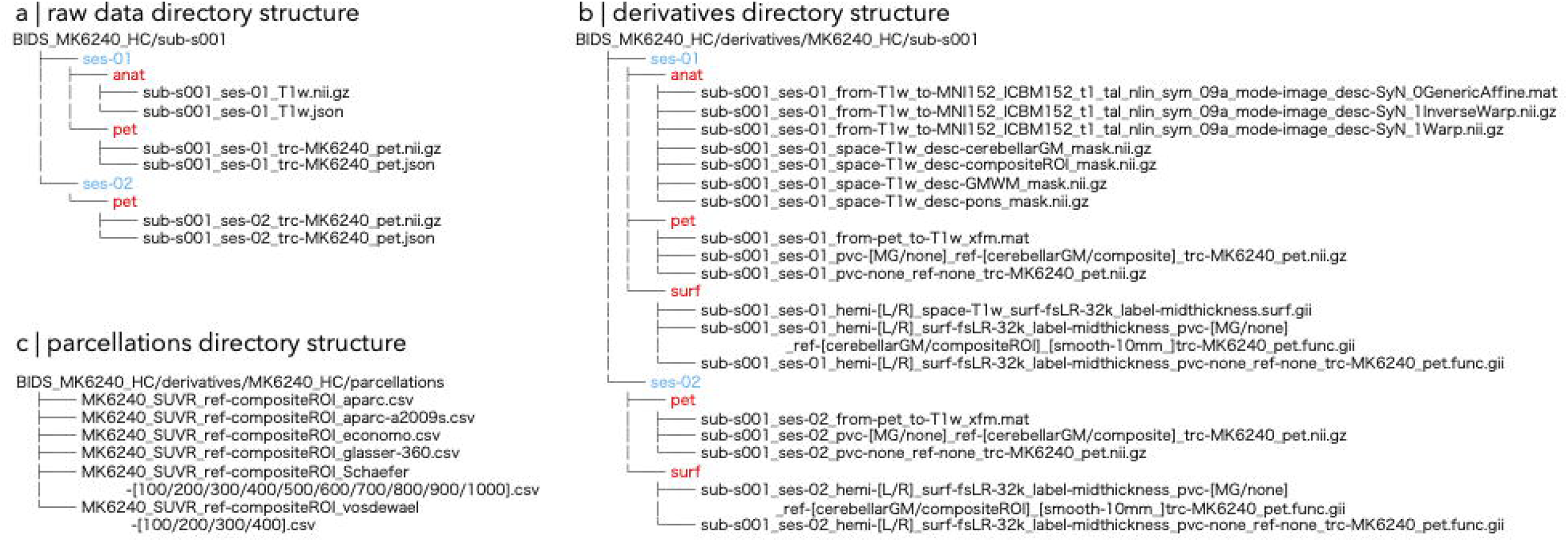
Directory structure of the [^18^F]MK6240 healthy control dataset for an example subject. (**a**) The *rawdata* directory contains anonymized, defaced T1w MRI (session 1) and 4D [^18^F]MK6240 PET images (session 1 and, when available, session 2). (**b**) The *anat* subdirectory of the *derivatives* directory contains the nonlinear registration files for the T1w MRI to MNI152 standard space, cerebellar grey matter and composite reference region masks, as well as a 4D grey matter and white matter mask input for partial volume correction with PVCPET. In the *pet* subdirectory, we include the linear registration file for the PET to T1w MRI, standard uptake value ratio (SUVR) volumes using the cerebellar grey matter and composite reference regions, both corrected and non-corrected for partial volume using Müller-Gärtner (MG) correction. A volume without normalization or partial volume correction corresponding to the 3D [^18^F]MK6240 image is also included. The *surf* subdirectory holds the individual fsLR32k midthickness surface as well as the previous volumes projected to fsLR32k space. Smoothed (10mm full-width at half-maximum) and non-smoothed maps are included. (**c**) Parcellated SUVR maps normalized to the composite ROI reference region in a number of parcellation schemes and resolutions are included as comma-separated value (csv) files.

### Derivatives

An affine registration file for the [^18^F]MK6240 PET to the T1w MRI, derived from ANTs, is provided in */derivatives/MK6240_HC/sub-#/ses-0#/pet*. In addition, non-linear registration files for the T1w MRI to MNI152 space can be found in */derivatives/MK6240_HC/sub-#/ses-0#/anat* (**Figure 2B**).

The cerebellar grey matter and composite ROI masks are contained in */derivatives/MK6240_HC/sub-#/ses-01/anat*. We also include an eroded pons mask for those who wish to use the pons as a reference region. Additionally, a two-frame image (*/derivatives/MK6240_HC/sub-#/ses-01/anat/sub-#_ses-01_space-T1w-desc-GMWM_mask*.*nii*.*gz*) containing probabilistic maps of grey matter and white matter, respectively, is included as input for the Müller-Gärtner partial volume correction in PETPVC toolbox.

Both Müller-Gärtner partial volume corrected and non-partial volume corrected data (pvc-MG and pvc-none) normalized to both cerebellar grey matter and composite reference regions (ref-cerebellarGM and ref-compositeROI) are included in */derivatives/MK6240_HC/sub-#/ses-0#/pet*. Additionally, the non-corrected and non-normalized PET image (*i*.*e*., the raw 4D [^18^F]MK6240 PET image averaged over time to a single 3D image) can be found as *sub-s#_ses-#_pvc-none_ref-none_trc-MK6240_pet*.*nii*.*gz*.

In the */derivatives/MK6240_HC/sub-#/ses-0#/surf* branch, we have included the individual mid-thickness surface file in fsLR32k resolution (*sub-s#_ses-01_hemi-#_space-T1w_surf-fsLR-32k_label-midthickness*.*surf*.*gii*) as well as surfaces derived from the five partial volume corrected and non-corrected and cerebellar grey matter, composite ROI normalized, and non-normalized volumes from the */pet* subfolder. Raw and smoothed (10mm FWHM) surface-mapped data are also provided.

The */derivatives/MK6240_HC/sub-#/ses-02* branch contains the same PET volumes and surfaces but for session-2. Again, no anatomically derived volumes and surfaces are provided in the session-2 subfolder as only one MRI session was available, and all session-2 PET data were registered to the session-1 MRI.

### Parcellations

Parcellated session 1 non-partial volume corrected [^18^F]MK6240 PET data normalized to the composite ROI reference region in comma separated value (csv) format are provided in the */derivatives/MK6240_HC/parcellations* folder. We include the following parcellation schemes: Desikan-Killany^49^ (aparc), Destrieux^50^ (aparc-a2009s), Von Economo,^51^ Schaefer^52^ (in a range of resolutions from 100-1000 nodes), Glasser^53^ (derived from the Human Connectome Project), and subparcellations of aparc^54^ (in a range of resolutions from 100-400).

### Technical Validation

#### Reference region and signal-to-noise

Previous studies have suggested that cerebellar grey matter is an ideal reference region for tau PET tracers due to its lack of involvement in neurodegenerative pathophysiology.^21,55,56^ One study examining [^18^F]MK6240 in AD patients and healthy controls both cross-sectionally and longitudinally showed that uptake in cerebellar grey matter remain similar across groups and across time.^56^ Another study on [^18^F]MK6240 comparing cerebellar grey matter and eroded white matter reference regions showed no advantage of eroded white matter over cerebellar grey matter and, in fact, it was prone to spill-over in AD patients.^32^ Landau and colleagues described a composite reference region consisting of the whole cerebellum, brainstem, and eroded subcortical white matter.^37^ They demonstrated that this approach enhanced sensitivity to longitudinal changes in beta-amyloid over time using florbetapir PET, compared to using the cerebellum or pons alone. Similar findings have been noted with tau tracers, showing an advantage to including supratentorial white matter for improved repeatability.^57–60^

Here, we included SUVR maps using either the cerebellar grey matter reference region or the composite reference region described by Landau and colleagues.^37^ The mean and standard deviation surface maps for non-corrected and Müller-Gärtner corrected PET data normalized to the cerebellar grey matter and composite reference regions are shown in **Figure 3A-B**. While [^18^F]MK6240 may show considerable spill over from meningeal uptake, our processing pipeline limits this effect by employing a partial volume correction technique, sampling to the midthickness surface, and spatially smoothing on the surface.

**Figure 3.**
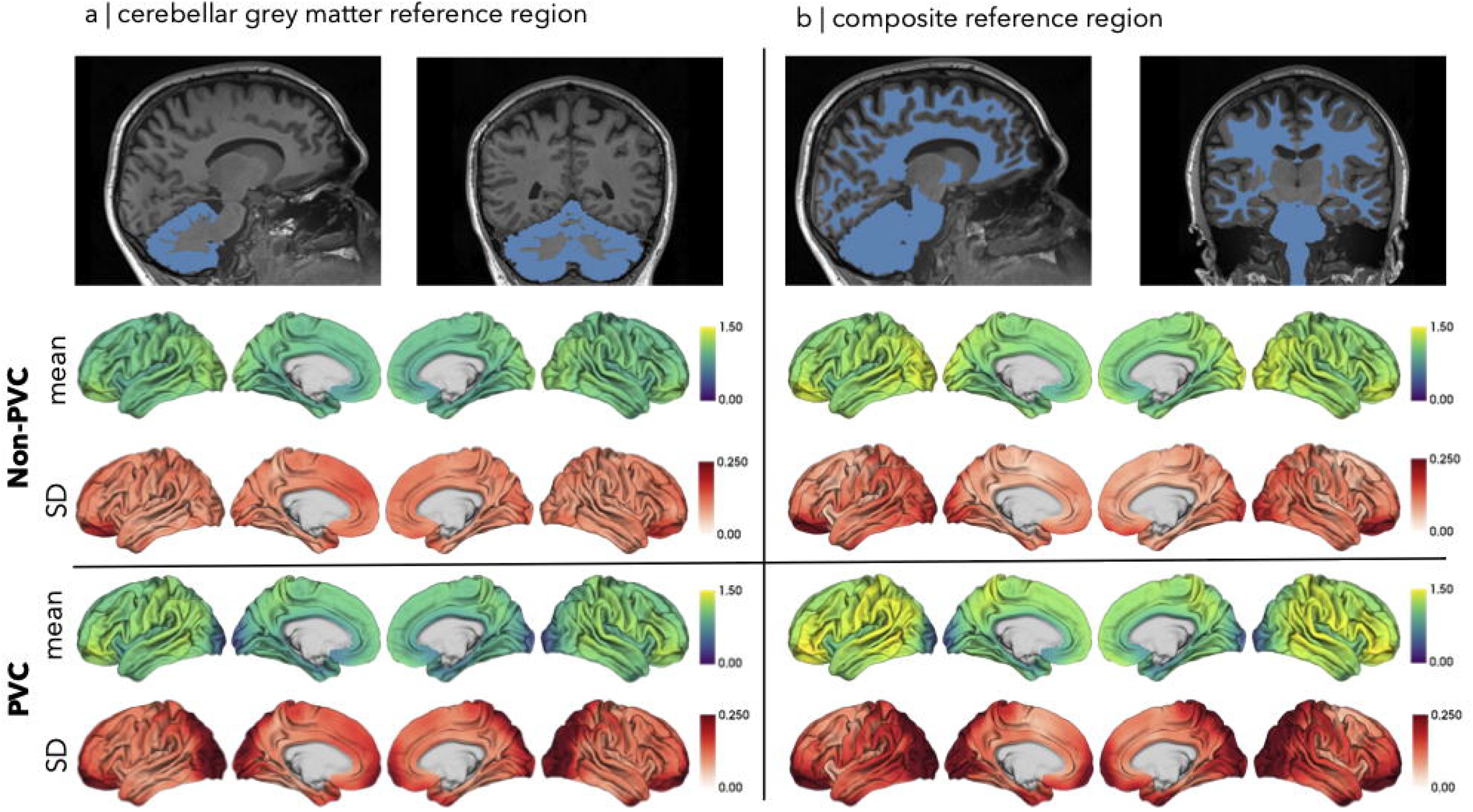
Mean standard uptake value ratio (SUVR) and standard deviation (SD) maps for (**a**) cerebellar grey matter and (**b**) composite ROI reference regions are shown. Partial volume corrected (PVC) data using the Müller-Gärtner method are shown in the bottom rows.

Ridgeline plots of the distribution of individual SUVRs for both cerebellar grey matter and composite ROI reference regions are shown in **Figure 4A-B**. Overall, the SUVRs skewed higher using the composite ROI reference region due to a smaller normalization factor. We calculated the signal to noise ratio (SNR) maps by dividing the raw PET maps by the average signal outside the head and then normalizing by the composite ROI reference region. We projected the SNR maps to the surface with the resulting average surface map shown in **Figure 4C**. There was generally high SNR, especially in inferior frontal, posterior temporal, and occipital areas.

**Figure 4.**
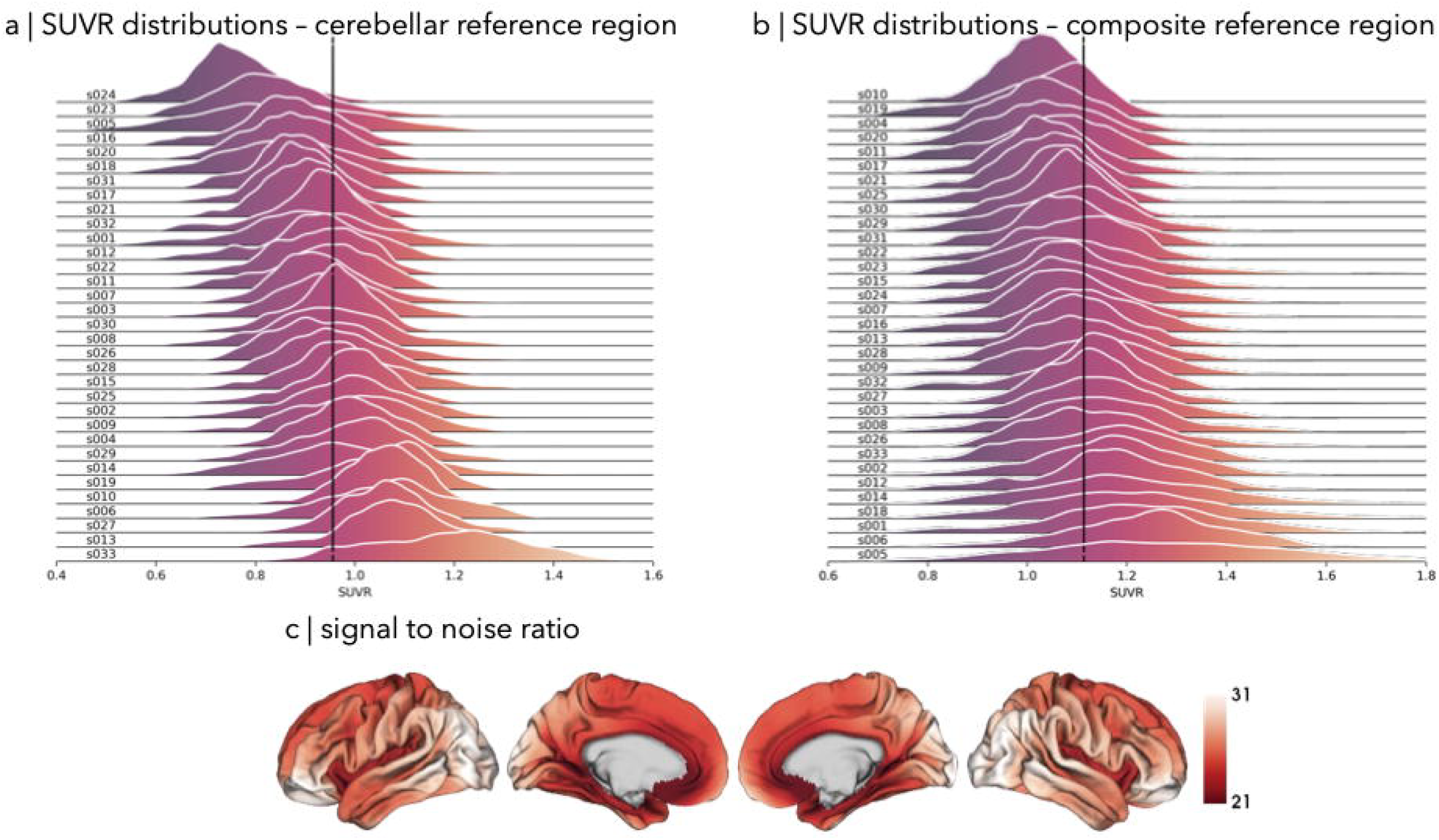
Ridgeline plots showing the distribution of standard uptake value ratio (SUVR) maps of individual participants are shown for the cerebellar grey matter (**a**) and composite (**b**) reference regions. The average signal to noise ratio (SNR) map is shown in (**c**) demonstrating high signal in occipital, posterior temporal, and inferior frontal areas.

### Intra- and inter-subject reliability and identifiability

Previous work has shown excellent 6-month test-retest reliability of [^18^F]MK6240 in healthy controls.^61^ In our subcohort of 9 individuals who had at least 1-year follow up [^18^F]MK6240 PET scan, we examined both the intra- and inter-subject reliability (session 1 *vs* session 1 and session 1 *vs* session 2) of partial volume corrected data with a composite ROI reference region. A correlation matrix shows the session 1 vs session 2 correlation for the same individual along the diagonal (intra-subject reliability) (**Figure 5A**). The bottom-left half of the correlation matrix represents the session 1 *vs* session 1 correlations between different individuals, while the upper-right half shows the session 1 *vs* session 2 correlations (inter-subject reliability). Mean intra-subject correlation across the 9 participants was 0.88±0.04 (**Figure 5B**), while the mean inter-subject correlation, calculated by averaging the correlations between subjects in session 1 *vs* session 1 and session 1 *vs* session 2, was 0.60±0.09. All individuals had test-retest correlation coefficients at or above 0.79.

**Figure 5.**
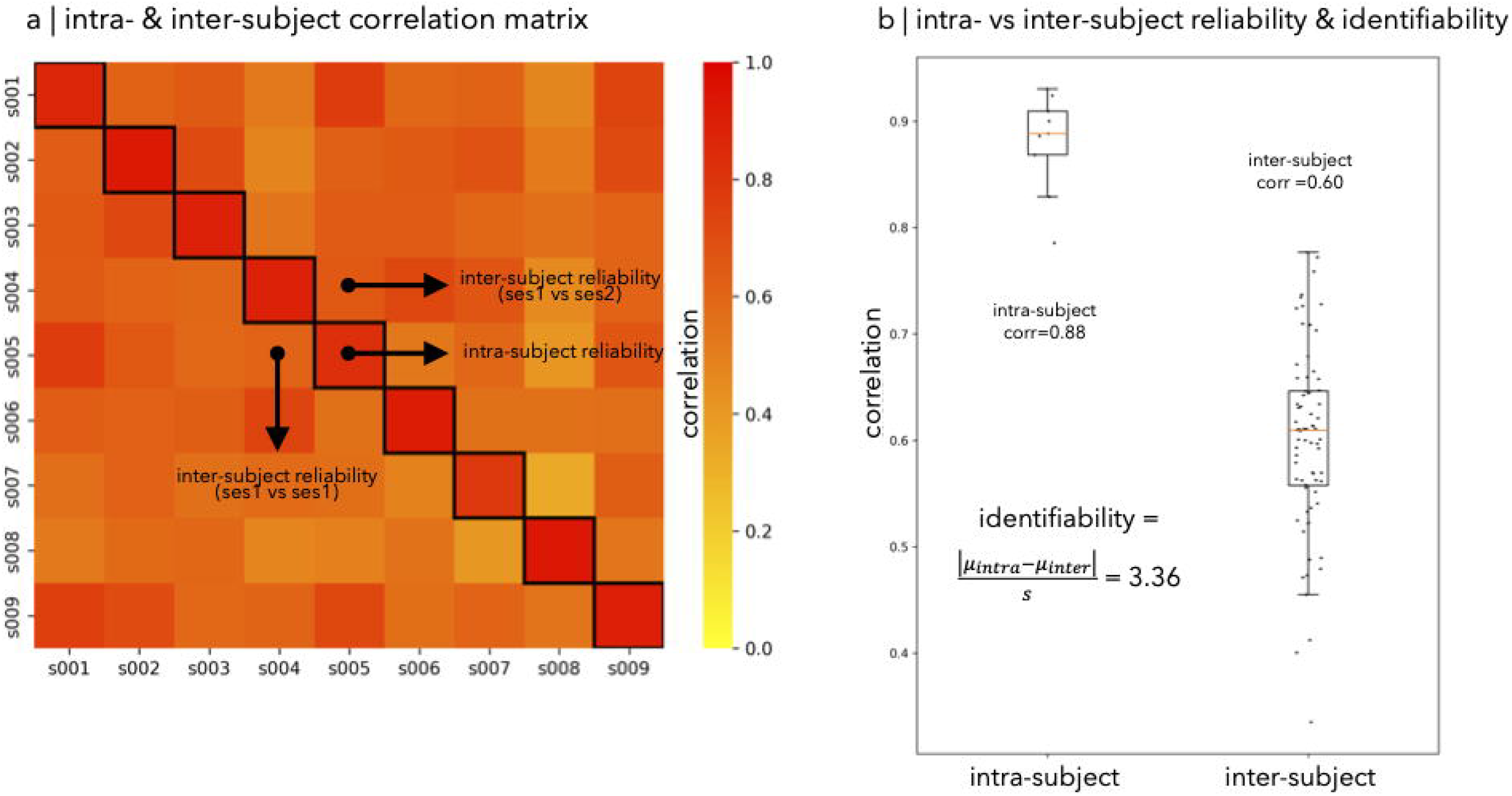
Correlation matrix of the nine subjects with retest data is shown in (**a**). Partial volume corrected data was used. The bottom-left half represents the inter-subject correlations for the session (ses) 1 scans while the top-right half represents the correlations between the session 1 vs session 2 scans. Outlined in black in the diagonal represents the intra-subject reliability, correlating each patient’s session 1 vs session 2 scan. Boxplots showing the median intra- and inter-subject correlations with whiskers extending to the minimum or maximum value within 1.5 times the interquartile range are shown in (**b**). The mean intra-subject (intra) and inter-subject (inter) correlations were 0.88 and 0.60, respectively. The identifiability was calculated to be 3.36.

Intraclass correlation was estimated based on a single-measurement, consistency, two-way mixed effects model using the Pingouin python statistical package.^62–64^ The intraclass correlation coefficient was 0.81, indicating good reliability.^63^

Next, we assessed how well [^18^F]MK6240 PET measures were reproducible across sessions while preserving individual differences by calculating the identifiability,^65–67^ which reflects the effect size of the difference between intra- and inter-subject reliability. This measure is calculated as the absolute difference in mean intra- and inter-subject correlations divided by the pooled standard deviation. The identifiability was calculated to be 3.36, which is comparable with previously reported neuroimaging biomarker, such as myelination,^66^ and 7T □microstructural profile covariance derived from quantitative T1 relaxometry and default mode network resting state functional MRI.^68^ Illustrative surface maps for the highest and the lowest correlating cases are displayed to the right of the boxplot (**Figure 6A**). Strength of test-retest correlation did not appear to be related to the length of interval between sessions (r = 0.22, p = 0.56). A vertex-wise correlation map is presented in **Figure 6B** showing generally good correlation across the cortex with slightly less correlation in inferior frontal, inferior temporal, and midline structures which may reflect some variability induced by meningeal uptake.

**Figure 6.**
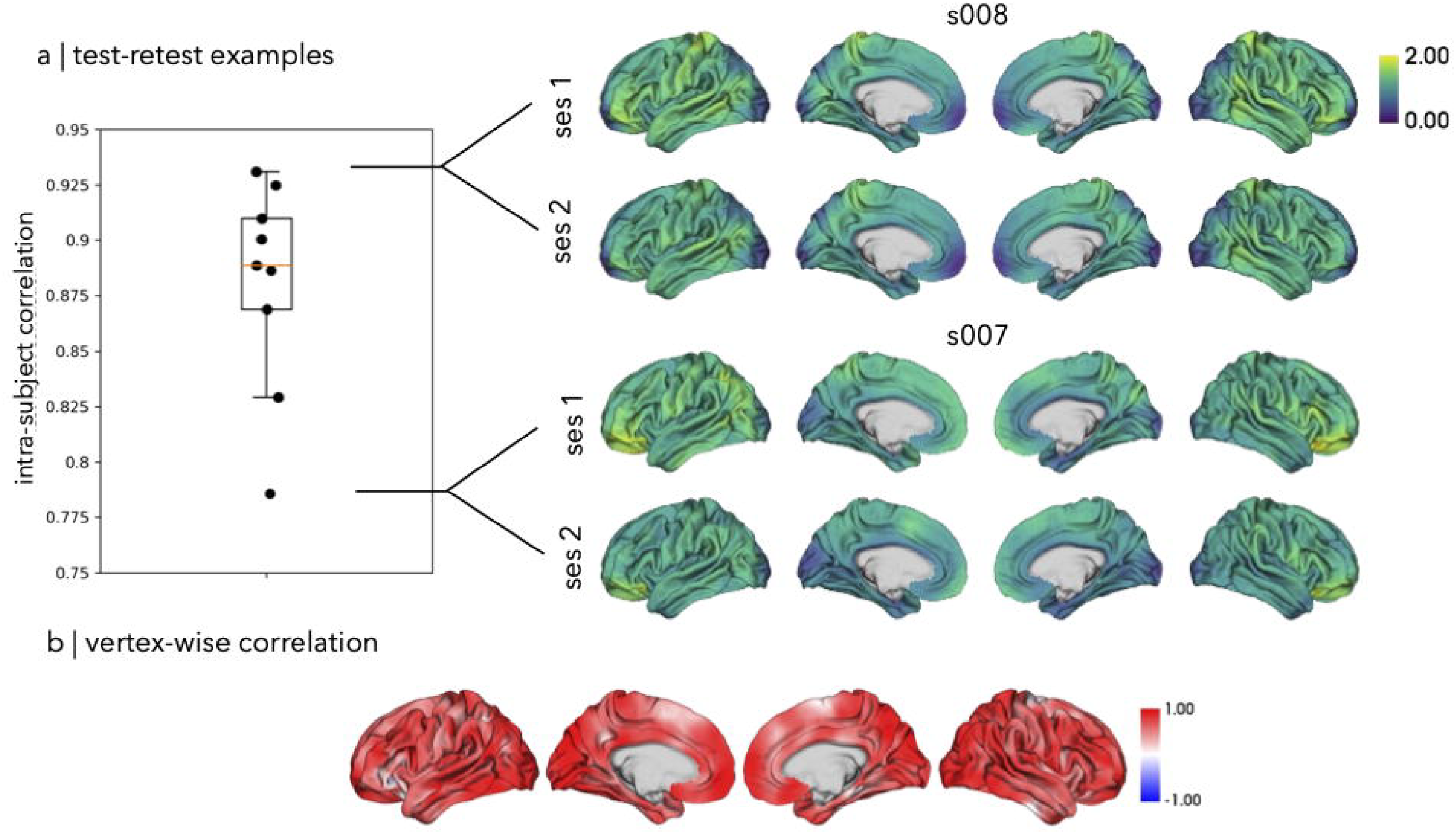
The intra-subject correlation boxplot from Figure 5b is shown zoomed up in (**a**) with the session 1 and session 2 [^18^F]MK6240 partial volume corrected maps of the highest and lowest correlated subjects (s008 and s007, respectively) shown to the right. Vertex-wise correlation was calculated across the 9 subjects in (**b**) showing generally good correlation across the cortex with slightly lower correlation in mesial temporal, anterior temporal and midline regions.

Similar analyses were performed on non-partial volume corrected data, which also showed good, albeit slightly lower, intra- and inter subject reliability (0.78±0.06 and 0.57±0.13, respectively) and identifiability (1.69). These data are presented in **Figures 7 and 8**.

**Figure 7.**
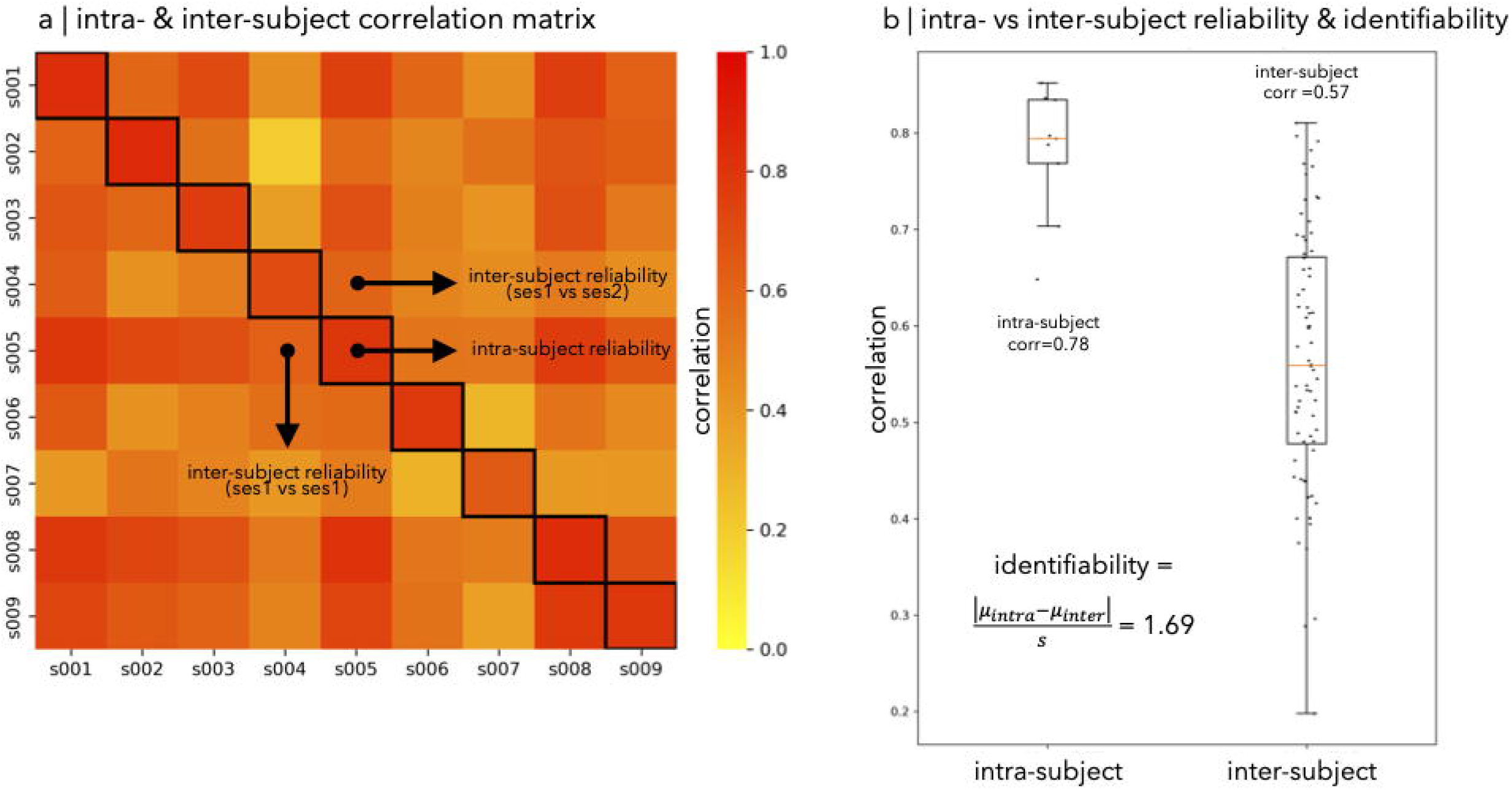
Correlation matrix of the nine subjects with retest data similar to Figure 5 except with non-partial volume corrected data. The bottom-left half represents the inter-subject correlations for the session (ses) 1 scans while the top-right half represents the correlations between the session 1 vs session 2 scans. Outlined in black in the diagonal represents the intra-subject reliability, correlating each patient’s session 1 vs session 2 scan. Boxplots showing the median intra- and inter-subject correlations with whiskers extending to the minimum or maximum value within 1.5 times the interquartile range are shown in (**b**). The mean intra-subject (intra) and inter-subject (inter) correlations were 0.78 and 0.57, respectively. The identifiability was calculated to be 1.69.

**Figure 8.**
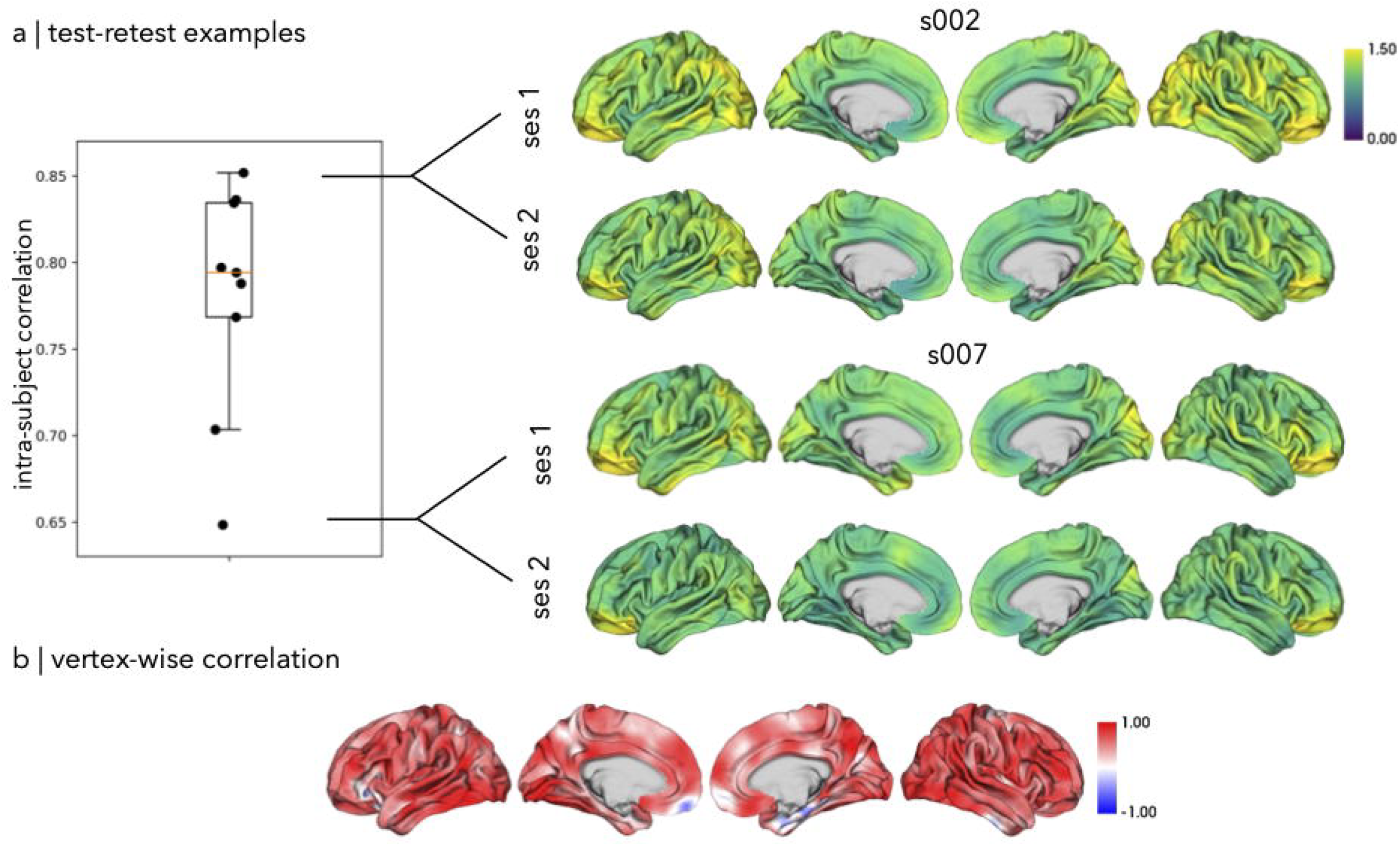
The intra-subject correlation boxplot from Figure 7 is shown zoomed up in (**a**) with the session 1 and session 2 [^18^F]MK6240 non-partial volume corrected maps of the highest and lowest correlated subjects (s002 and s007, respectively) shown to the right. Vertex-wise correlation was calculated across the 9 subjects in (**b**) showing generally good correlation across the cortex with poor/negative correlation in mesial temporal, inferior frontal, and inferior temporal areas.

Overall, in this small subcohort of participants with a follow up scan, [^18^F]MK6240 shows good identifiability and test-retest reliability.

## Data availability

This dataset is available for download on the Open Science Framework (https://doi.org/10.17605/OSF.IO/ZNT9D).^48^

## Code availability

The T1w MRI processing pipeline, micapipe v0.2.3, is openly available on GitHub (https://github.com/MICA-MNI/micapipe) and extensively documented on ReadTheDocs (https://micapipe.readthedocs.io/). PET partial volume correction was done using the PETPVC toolbox (https://github.com/UCL/PETPVC). Code for figure generation and derivatives generation are available on our GitHub (https://github.com/jlam-nsx/MK6240_HC_data_release).

## Acknowledgements

We would like to thank the participants for their efforts to take part in the study and to be willing to contribute to the Montreal Neurological Institute’s open science initiative. We also thank the members of the McConnell Brain Imaging Centre’s MRI and PET units for their help in data acquisition and processing. JL is funded by the Fonds de la Recherche du Québec – Santé (FRQS) and the Ministère de la Santé et des Services sociaux du Québec (MSSS). RRC received support from the FRQS, the Montreal Neurological Institute Jeanne Timmins Costello Fellowship, and the Healthy Brains, Healthy Lives – Entrepreneur Postdoctoral Fellowship. JR received support from the Canadian Open Neuroscience Platform (CONP) and Canadian Institutes of Health Research (CIHR). LC acknowledges prior support from Brain Research UK (Award 14181). BCB acknowledges support from CIHR (FDN-154298, PJT-174995, PJT-191853), SickKids Foundation (NI17-039), Natural Sciences and Engineering Research Council (NSERC RGPIN-2025-05932), Azrieli Center for Autism Research of the Montreal Neurological Institute (ACAR), BrainCanada, FRQS, the Helmholtz International BigBrain Analytics and Learning Laboratory (HIBALL), the Canada Research Chairs Program (CRC), and the Centre of Excellence in Epilepsy at the Neuro (CEEN).

## Author contributions

Study design and conception: JL, LC, MK, AB, NB, BCB; data acquisition, analysis, interpretation: JL, RRC, TA, JR, JC, AD, ES, RH, CH, GM, BCB; manuscript drafting and data organization: JL, RRC, BCB. All authors provided feedback and approved of the final manuscript.

